# A chicken IgY can efficiently inhibit the entry and replication of SARS-CoV-2 by targeting the ACE2 binding domain in vitro

**DOI:** 10.1101/2021.02.16.430255

**Authors:** Jingchen Wei, Yunfei Lu, Ying Rui, Xuanyu Zhu, Songqing He, Shuwen Wu, Qing Xu

## Abstract

COVID-19 pneumonia has now spread widely in the world. Currently, no specific antiviral drugs are available. The vaccine is the most effective way to control the epidemic. Passive immune antibodies are also an effective method to prevent and cure COVID-19 pneumonia. We used the SARS-CoV-2 S receptor-binding domain (RBD) as an antigen to immunize layers in order to extract, separate, and purify SARS-CoV-2-IgY from egg yolk. SARS-CoV-2-IgY (S-IgY)can block the entry of SARS-CoV-2 into the Cells and reduce the viral load in cells. The Half effective concentration (EC_50_) of W3-IgY (S-IgY in the third week after immunization) is 1.35 ± 0.15nM. The EC_50_ of W9-IgY (S-IgY in the ninth week after immunization) is 2.76 ± 1.54 nM. When the dose of S-IgY is 55 nM, the fluorescence representing intracellular viral protein is obviously weakened in Immunofluorescence microscopy.

Results of Sars-CoV-2 /Vero E6 cell experiment confirmed that S-IgY has a strong antiviral effect on SARS-CoV-2, and its (EC_50_) is 27.78 ±1.54 nM*vs* 3,259 ± 159.62 nM of Redesivir (differ > 106 times *P<0*.*001*).

S-IgY can inhibit the entry and replication of SARS-CoV-2, which is related to its targeting the ACE2 binding domain.

S-IgY is safe, efficient, stable, and easy to obtain. This antibody can be an effective tool for preventing and treating COVID-19 pneumonia.

Fig. 1.
Graphical Abstract
The figure briefly illustrates that the preparation and extraction of S-IgY and its anti-S-CoV-2 mechanism is to inhibit the entry and replication of SARS-CoV-2 by targeting the ACE2.

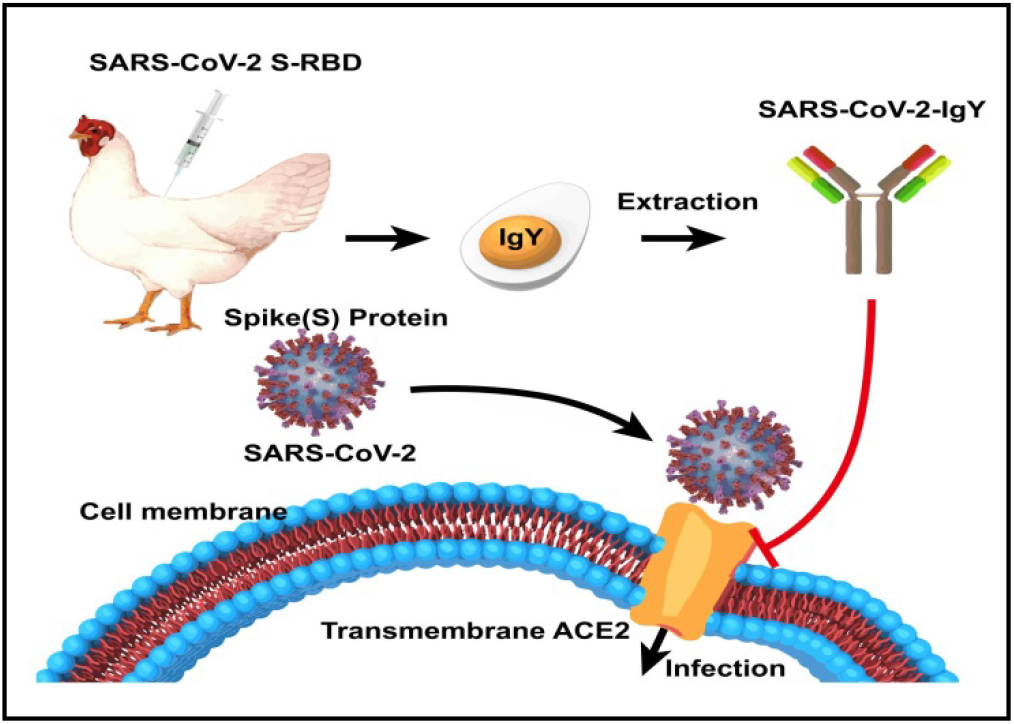

## Introduction

The 2019 novel coronavirus (SARS-CoV-2) is a new coronavirus strain found in the human body in 2019. During the incubation period, the virus can be transmitted. The coronavirus’ main transmission routes are air transmission and contact transmission, and the intermediate host is still unclear (1). As a global “gray rhinoceros” and “black swan event”, the COVID-19 pandemic impacts human health and influences the world’s economy and politics. The severity of impact depends on when the pandemic started to spread and the intensity of prevention and control measures in various countries (2) (source: WHO). By modeling and analyzing factors such as the season, mode of transmission, and cross-immunity between immunity and coronavirus, it is predicted that the COVID-19 pandemic may last until 2024(3).

SARS-CoV-2 can be divided into three categories: A, B, and C. Type A is the root of the virus outbreak, concentrated in infected people in the United States and Australia. B strains are mutated from A, mainly in China (i.e., Wuhan). Type C evolved from type B, which infected people in Europe primarily carry (4). At present, many SARS-CoV-2 mutants have spread in Britain, France, India, Denmark, Japan, Portugal, Norway, Jordan, South Korea, and other places, making SARS-CoV-2 more contagious and more difficult to prevent and control (source: WHO)(5).

The spike protein S, envelope E, membrane glycoprotein M, and nucleocapsid N are the four essential coding structural proteins for coronavirus to complete viralassembly and infection. It was found that the S protein located on the surface of the SARS-CoV-2 virus, is the fusion protein of the virus. The S protein is responsible for combining the virus and the host cell receptor, which mediates the first step of SARS-CoV-2 infection (6). Each trimeric S protein monomer is about 180 KD and contains two subunits, S1 and S2, mediating attachment and membrane fusion. In the structure, the N- and C-terminal portions of S1 fold as two independent domains, the N-terminal domain (NTD) and C-terminal domain (C-domain). Depending on the virus, either the NTD or C-domain can serve as the receptor-binding domain (RBD) (7, 8). The receptor of SARS-CoV-2 invading human cells is angiotensin-converting enzyme 2(ACE2) (9).

The primary pathological process of patients with COVID-19 is the inflammatory reaction caused by the SARS-CoV-2 virus combined with ACE2. Fever, dry cough, and fatigue are common symptoms of COVID-19. Most patients have bilateral frosted glass shadow changes and lymphocyte reduction in chest CT scan. Patients with underlying medical conditions and older individuals are more likely to have severe illness and death after infection than healthier and younger individuals (10, 11).

The results of a large-scale randomized controlled trial (RCT) of glucocorticoids for treatingCOVID-19 pneumonia showed that short-term oral or intravenous administration of low-dose hormone could greatly reduce the mortality of patients who need oxygen inhalation or a non-invasive ventilator within 28 days (12). The U.S. FDA approved Remdesivir as the first COVID-19 treatment drug in the United States. Remdesivir might shorten the time to clinical improvement among hospitalized adults with severe COVID-19 (13). However, it was not recommended as a first-line treatment by the World Health Organization (14). Since the COVID-19 outbreak, countries around the world have accelerated the development of the COVID-19 vaccine. By December 2020, 60 candidate COVID-19 vaccines have been approved for clinical trials, and seven of these vaccines (three inactivated vaccines, two mRNA nucleic acid vaccines, and two vector vaccines) have been approved for emergency use or conditional marketing (source: WHO) (15,16). The effective rate of the mRNA vaccine of Pfizer pharmaceutical co., ltd. in the United States is 95%. The production and persistence of immune neutralizing antibodies in vaccinated people need to be confirmed (17).

Antibodies that can effectively neutralize SARS-CoV-2 were found in RBD-specific monoclonal antibodies from B lymphocytes of patients. This activity is related to the competition between the antibody and ACE2 for binding RBD (18). Using a recombinant human ACE2 antibody protein purified in vitro to “neutralize” the SARS-CoV-2 virus can reduce its infection ability by 1,000-5,000 times (19). The mechanism of plasma therapy is to use the specific antibody against the virus found in the recovered patients’ plasma to quickly identify and capture and eliminate the virus through the activation of the body’s complement system. Plasma therapy has achieved good results in major epidemics such as SARS, MERS, Ebola, and H1N1 influenza (20). Recent research data showed that the transfusion of the plasma that contains high titer neutralizing antibodies from the SARS-CoV-2 infected patients in convalescence to the critically ill patients can restore the organ functioning within a short time, give a negative result in virus nucleic acid detection and increase specific antibody concentration (21). However, because of the shortage of plasma in convalescence, the disparity between plasma demand and supply, and donor plasma safety, it is still not feasible to obtain qualified plasma in convalescence.

Egg yolk Immunoglobulin Y (IgY) extracted from immunized poultry egg yolk is an excellent source of the antibody for passive immunization (22). IgY has long been used to prevent and treat infectious diseases of poultry and livestock (23), and research on IgY in the diagnosis and treatment of human diseases has become increasingly popular. For example, anti-*Helicobacter pylori* (24), Shiga *Escherichia coli* (25), and *Vibriocholerae*(26), as well as other bacteria, anti-human rotavirus (27), influenza virus (28,29), SARS virus (30), and the hand, foot, and mouth disease (31), all used IgY as a treatment method. Here, we report an egg antibody (SARS-CoV-2-IgY) study with a high neutralization ability against SARS-CoV-2 in vitro.

## Results

### 1. The expression and purification of SARS-CoV-2 S-RBD

SDS-PAGE results showed that the fusion protein of SARS-CoV-2 S-RBD was between 70kD and 55kD, in according with the predicted molecular weight 66.2KD was made by the pCMV-DsbC-RBD fusion expression vector (provided by Sina Biological Company) and band one and two showed the purity of the protein was very high (Fig.2).

**Fig. 2.**
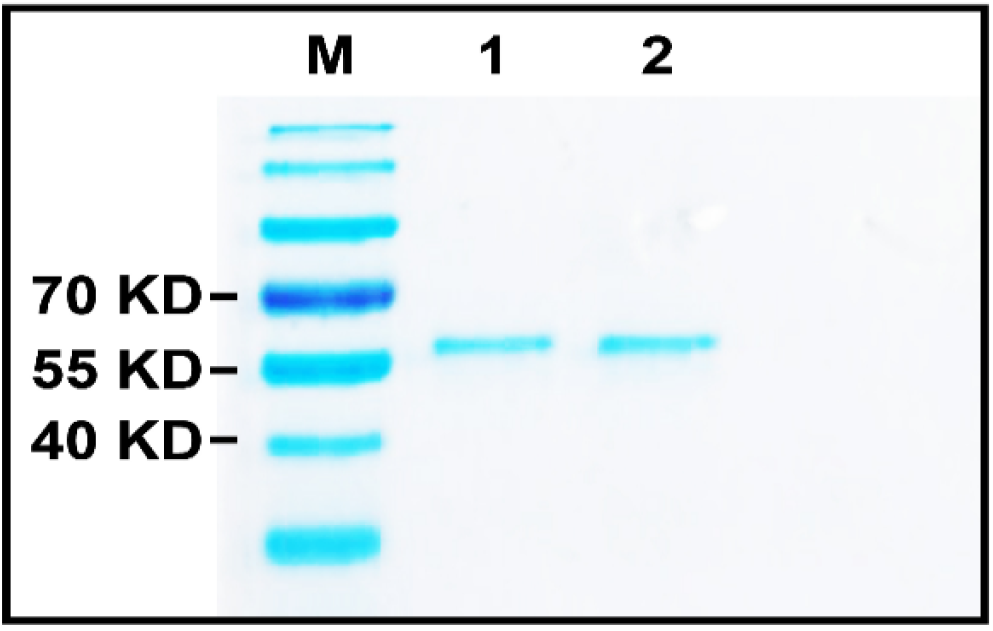
Gel electrophoresis of SARS-CoV-2-S-RBD. Fusion protein of SARS-CoV-2 S-RBD was between 70kD and 55kD, in according with the predicted molecular weight 66.2KD was made by the pCMV-DsbC-RBD fusion expression vector (provided by Sina Biological Company) and band one and two showed the purity of the protein was very high.

### 2. The preparation of total IgY

The laying hens were immunized with antigen of SARS-CoV-2 S-RBD for 4 times, we obtained total S-IgY against SARS-CoV-2 from egg yolk by ways of water extraction, salting out, dialysis and ultrafiltration, etc. Results of the gel electrophoresis SDS-PAGE showed that the heavy chain band was at 70kD and the light chain band was at 25kD, and the total IgY purity was about 85% (Fig.3).

**Fig. 3.**
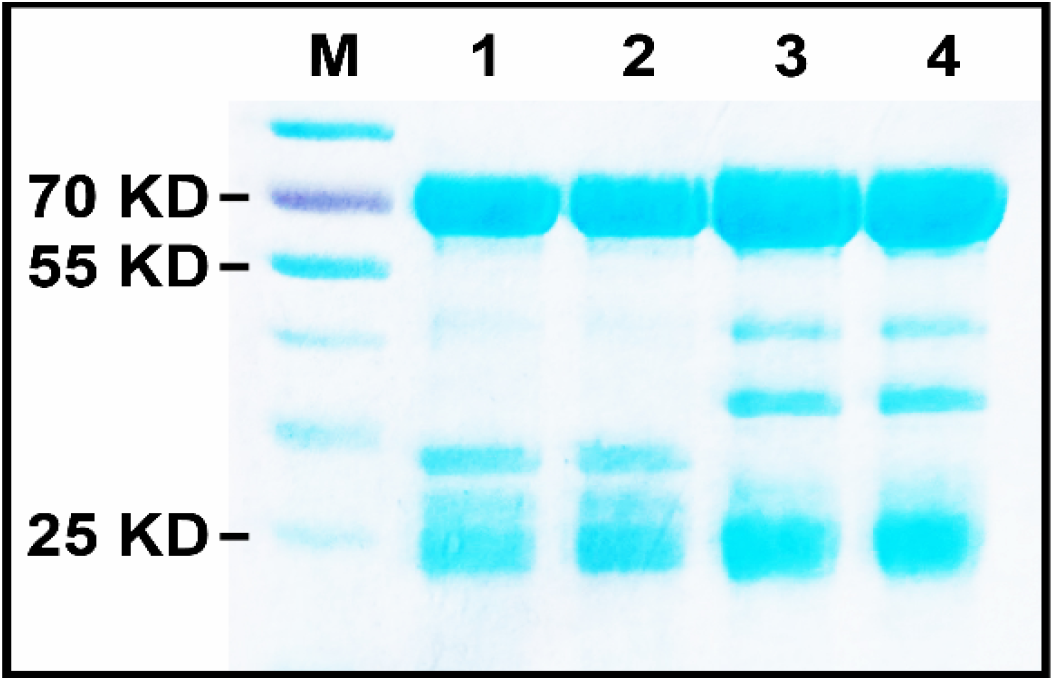
SDS-PAGE gel detection of S-IgY. SDS-PAGE gel detection of S-IgY was done which showed that the heavy chain band was at 70kD and the light chain band was at 25kD, and the total IgY purity was about 85%.

### 3. The specificity of S-IgY

In order to confirm the recognition and binding ability of SARS-CoV-2 to S-RBD antigen, Western blot were done. The dilution of primary antibody (S-IgY) was 1:10000, and the dilution of secondary antibody (HRP-labeled goat anti-chicken) was 1: 10000. The results showed that S-IgY had a good effect on S-RBD antigen The antibody band was clearly located between 55 KD and 70KD, which was consistent with the electrophoresis position of S-RBD antigen (66.2KD), and there were few hetero bands, which indicated that the content of nonspecific IgY in total IgY was less, and the proportion of specific S-IgY was higher. (Fig.4).

**Fig. 4.**
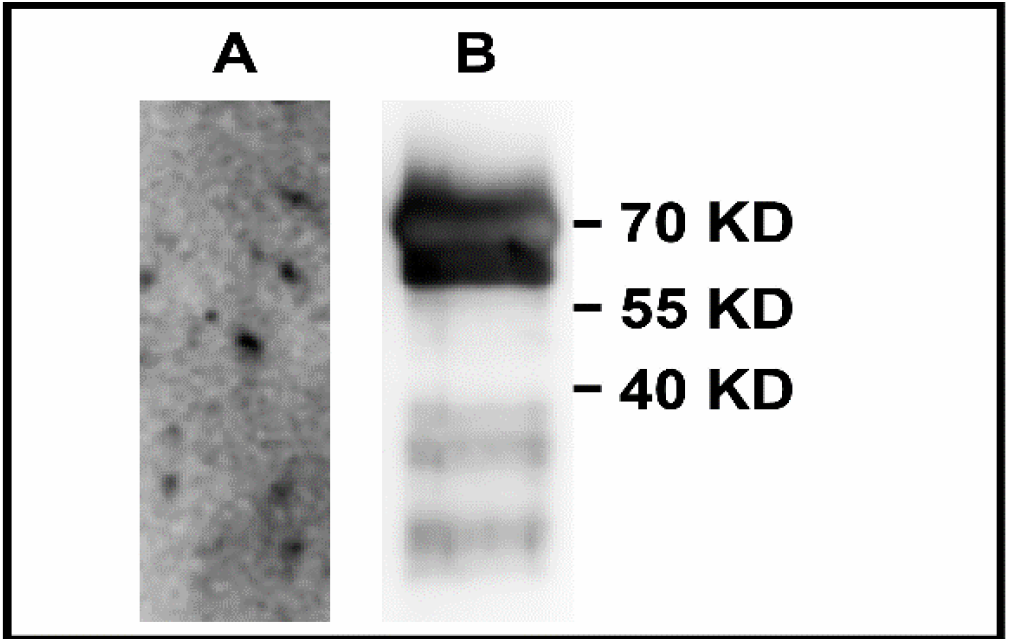
Detection of SARS-CoV-2-IgY specificity by Western blot. **A:** Pre-Immu-IgY **B:** SARS-CoV-2–IgY(66.2KD) The WB results showed that SARS-CoV-2-IgY(S-IgY) had a good effect on S-RBD antigen The antibody band was clearly located between 55 KD and 70KD, which was consistent with the electrophoresis position of S-RBD antigen(66.2KD).In this experiment, the dilution of primary antibody (S-IgY) was 1:10000, and the dilution of secondary antibody (HRP-labeled goat anti-chicken) was 1: 10000.

### 4. S-IgY blocks cell entry of SARS-CoV-2

For investigating the role of S-IgY in neutralizing SARS-CoV-2 and preventing SARS-CoV-2 from entering target cells, Pseudo typed VSV luciferase-reporter particles bearing SARS-CoV-2 spike (S) protein (pSARS-CoV-2) were used to reflect the virus entry activity was done. The results of viral load in the cells detected by real time qPCR demonstrated that S-IgY could efficiently blocking the cell entry of virus. The EC_50_ of W3-IgY is 1.35 ± 0.15 nM and EC_50_ of W9-IgY is 2.76 ± 1.54 nM (Fig.5).

**Fig. 5.**
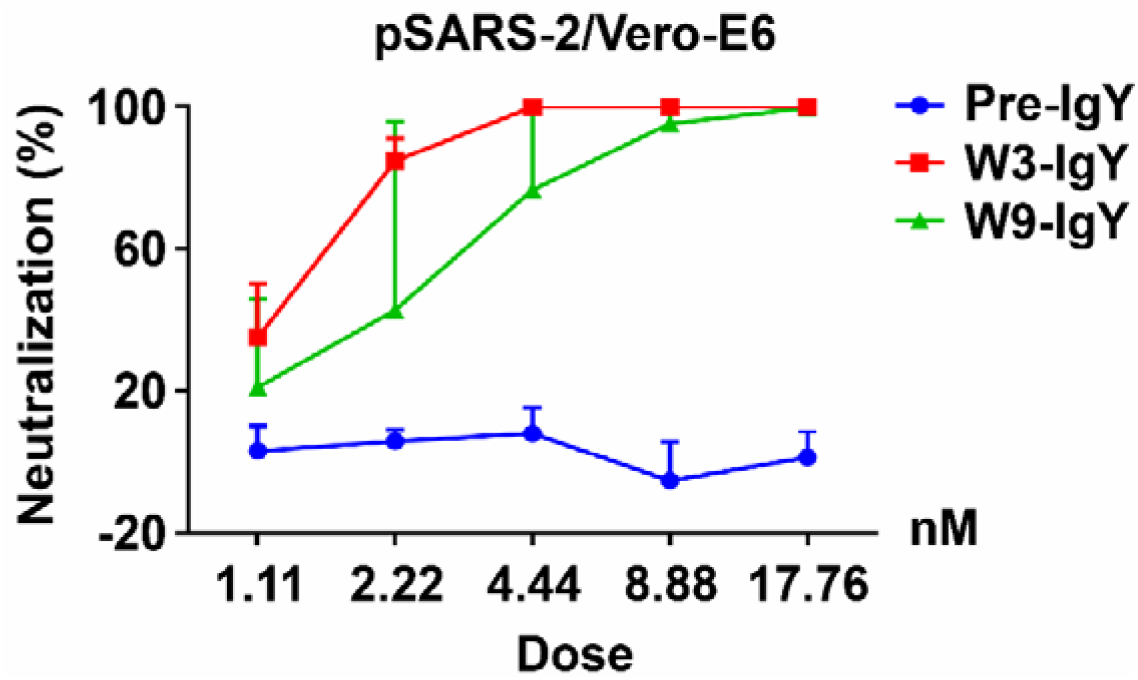
pSARS-CoV-2/VeroE6 (qRT-PCR) The resultsof viral load in the cells detected by qRT-PCR demonstrated that S-IgY can efficiently blocking the cell entry of virus, The EC_50_ of W3-IgY is 1.35 ± 0.15 nM and EC_50_ of W9-IgYis 2.76 ± 1.54 nM, and the Pre-IgY has no inhibitory effect on virus.

### 5. S-IgY reduce the replication of SARS-CoV-2

#### 5.1 Immunofluorescence

Cell Immunofluorescence experiment was done to detect the SARS-CoV-2-S protein expression in VeroE6cells, the results showed that that S-IgY could effectively inhibit the expression of SARS-CoV-2-S protein. SARS-CoV-2-S protein were observed in the pre-immunized IgY (Pre-IgY) group. When the concentration of S-2-IgY was 55nM, the SARS-CoV-2-S protein fluorescence in the field of vision was significantly reduced. About 50% of the fluorescence was suppressed; as the concentration of S-IgY was 110 nM, the fluorescence disappeared completely in the field of vision. The SARS-CoV-2 was completely suppressed by S-IgY (Fig.6).

**Fig. 6.**
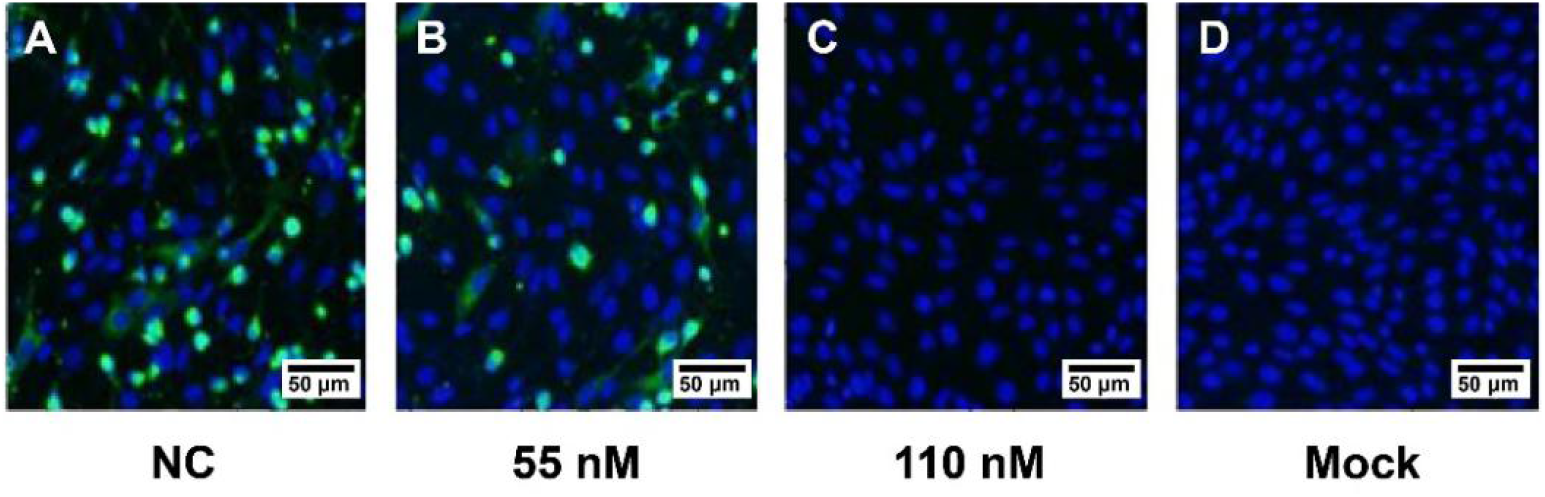
Immunofluorescence microscopy. Vero E6 cells were incubated with the S-IgY for 2 h, followed by incubation with the secondary antibody [Alexa 488-labeled goat anti-mouse (1:500; Abcam)]. The nuclei were stained with DAPI dye (Beyotime, China). The images were taken by fluorescence microscopy, Results were as follow: (A) NC:pre-immunizedIgY: Full-field virus protein (B) S-IgY 55 nM:Viral protein decreased significantly in visual field,About 50% of the viruses were suppressed; (C) S-IgY 110 nM:Virus protein disappeared completely in visual field, The virus was completely suppressed; (D) Mock Cells without virus addition. It means that S-IgY can effectively inhibit the SARS-CoV-2 from entering cells and has a dose-dependent relationship.

#### 5.2 qRT-PCR

SARS-CoV-2 (100PFU) was incubated with an equal volume of Pre-IgY, S-IgY and Remdesivir in different concentration gradients in Vero E6 cells. The results showed that different doses of S-IgY had an obvious inhibitory effect on the cellular SARS-CoV-2 content (copy number) with EC_50_ of 27.78 ± 1.54 nM. Remdesivir’s EC_50_ against SARS-CoV-2 was 3,259 ± 159.62 nM,

The inhibitory effect of S-IgY on SARS-Cov-2 replication was 106 times stronger more than that of Remdesivir In vitro (Fig.7).

**Fig. 7.**
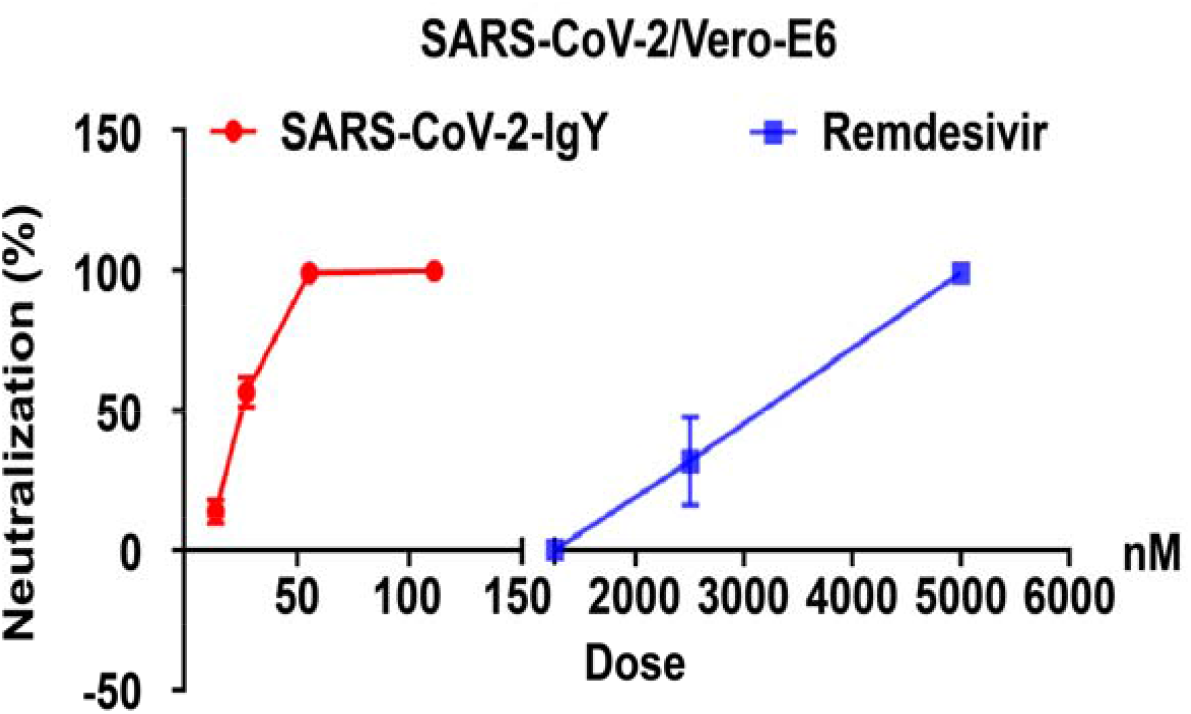
SARS-CoV-2/Vero-E6 (qRT-PCR) 25μL of SARS-CoV-2 (100PFU) was incubated with an equal volume of Pre-IgY, S-IgY and Remdesivir in different concentration gradients for 10 min, then added to 1×10^4^vero cells, and cultured in an incubator at 37°C for 4 h, and then the cell culture solution was changed to complete medium. After 48 h, cells were collected and total RNA was extracted, the copy number of SARS-CoV-2 genome in each sample was detected by qRT-PCR, EC_50_ of S-IgY is 27.78 ± 1.54 nM, and EC_50_ of Remdesivir is 3,259 ± 159.62 nM, S-IgY *vs* Remdesivir (*P*< 0.001). In vitro, the anti-SARS-CoV-2 effect of S-IgY is 106 times stronger than that of Remdesivir.

## Discussion

Since the outbreak of infectious diseases of SARS-CoV-2 in early 2019, COVID-19 virus has quickly spread worldwide. Currently, the World Health Organization believes that no specific drug can produce antiviral effects except the use of glucocorticoid for severe patients. The main hope of controlling the COVID-19 pandemic is placed on the use of vaccines. However, the vaccine also has some problems, such as vaccine failed due to unsuitability for people, short supply, individual differences in immune response, antibody titer and retention time, etc. Although keeping social distance including the use of face masks is an effective way to prevent infection, common surgical masks cannot completely block the air transmission of SARS-CoV-2 (30). Furthermore, preventive medicine and external disinfectants cannot be used in real-time. To sum up, it is very necessary to develop new control methods for SARS-CoV-2.

There is convincing scientific evidence for the prevention and treatment of passive immune specific antibodies in SARS-CoV-2. However, human antibodies are expensive and need to be strict cryopreservation which are not suitable as alarge-scale popularization and application.

We used SARS-CoV-2 S-RBD as an antigen to immunize hens to obtain SARS-CoV-2-IgY. Our study results confirmed that this antibody could recognize SARS-CoV-2 S-RBD antigen and specifically bind with it, thereby blocking the interaction between the S protein and the ACE2 and preventing infection caused by the interaction between the S protein and ACE2. The results of Real-time quantitative RT-PCR(qRT-PCR)and the immunofluorescence experiment have proved that S-IgY can prevent SARS-CoV-2 from entering human cells.

Resuts of SARS-CoV-2 /Vero E6 cell experiment confirmed that S-IgY has strong antiviral effect on SARS-CoV-2, and its EC_50_ was 27.78 ± 1.54 nM*vs* 3,259± 159.62 nM of Redesivir (differ > 106 times, *P*<0.001).

S-IgY can not only block SARS-CoV-2 from entering target cells, but also effectively inhibit replication of SARS-CoV-2 in cells.

For critically ill patients with pneumonia in COVID-19, the inflammatory storm caused by antibody therapy is also a problem worthy of great attention

From a study among patients in SARS-CoV-2, Wuhan, China in 2019, ICU patients had higher plasma levels of IL-2, IL-7, IL-10, GSCF, IP10, MCP1, MIP1A, and TNFα compared to non-ICU patients (31). Hypersensitivity inflammatory storm caused by viral infection is considered to be associated with the activation of complement C5a (32). IgY does not activate complement, so it is safer than other antibodies. IgY has more stable physical chemical properties and heat resistance than IgG: It is stable at 60°C-70°C and stable at 4°C for five years, at room temperature for 6 months (33, 34), It is easy to collect (35) and high in content. Each egg contains about 1.0 mg of specific antibody, which is equivalent to the content of the whole blood of a rabbit (36). Further, it is Acid-and enzyme-resistant and can be administered orally (37, 38). Its role in the prevention and treatment of SARS-CoV-2 infection deserves further study.

## Methods

### 1. Preparation of antigen

An enterokinase site sequence (DDDDK) was added to the N-terminal of the RBD peptide, according to prokaryotic expression preferences and the corresponding nucleic acid sequence was synthesized with reference to prokaryotic expression preference. The pCMV-DsbC-RBD fusion expression vector was constructed, and the DsbC-RBD fusion protein was obtained from eukaryotic expression and affinity purification. The fusion protein was digested with enterokinase and purified by secondary affinity to obtain a high purity SARS-CoV-2 S protein RBD peptide (provided by Sina Biological Company) (Fig.2).

### 2. Immunity of hens

SPF 14-week-old white single-crowned Leghorn hens (provided by SPF Experimental Animal Center of Guangdong XinxingDahua Agricultural Poultry Eggs Co., Ltd.) were reared at 20°C-25°C, 60%-90% humidity, natural light, and 100-level air purification. The food and water were disinfected. After four weeks of adaptive rearing, two weeks after laying eggs, the hens began to be immunized. The RBD protein antigen was mixed with the same amount of Freund’s incomplete adjuvant and fully emulsified for later use (the amount of antigen was 50-200 μg). Intramuscular injections of 0.25 mL were given to the armpit of bilateral chicken wings and the left and right sides of the abdomen. Immunization was performed once every seven days for a total of four times.

### 3. Extraction and separation of IgY

The egg yolk was separated with an egg separator, diluted nine times in water, and stirred well. The egg liquid’s pH was adjusted to 5.0-5.2 with 0.1 mol/L HCl, and then incubated overnight at 4°C. The samples were then frozen and centrifuged at 4,000 rpm for 40 min. (NH4)_2_SO_4_ was added to the supernatant to achieve a final saturation of 45% and incubated at 4°C for 3 h, followed by another freezing and centrifugation step at 4,000 rpm for 10 min. The supernatant was discarded, and water was added to dissolve the protein precipitate. Na_2_SO_4_ (final mass fraction of 13%) was added to dissolve the sample, followed by incubation at 4°C for 3 h. The samples were frozen and centrifuged at 4,000 rpm for 10 min, the supernatant was discarded, and PBS was added for dissolution. The solution was dialyzed in a dialysis bag for 4-5 h and stored at −20°C.

Separated egg yolk with egg separator, diluted egg yolk with 9 times of water, stirred well, adjusted pH of egg liquid to 5.0-5.2 with 0.1 mol/L HCl, and then stood at 4°C overnight. Freeze-centrifuge at 4000rpm for 40min, added (NH4)_2_SO_4_ to the supernatant to make its final saturation 45%, and stood at 4°C for 3h; freeze-centrifuge at 4,000 rpm for 10 min, centrifuge at 4,000 rpm for 10 min, discarded the supernatant, added water to dissolve the protein precipitate, added Na_2_SO_4_ (the final mass fraction is 13%) to fully dissolve it, and then stood at 4°C for 3h; freeze-centrifuge at 4,000 rpm for 10min, discarded the supernatant, and added PBS for dissolution. Dialyzed the solution in a dialysis bag for 4-5 h and stored it at −20°C.

Separated egg yolk with egg separator, dilute egg yolk with 9 times of water, stir well, adjusted pH of egg liquid to 5.0-5.2 with 0.1 mol/L HCl, and then stand at 4°C overnight. Freeze-centrifuge at 4,000 rpm for 40 min, added (NH_4_)_2_SO_4_ to the supernatant to make its final saturation 45%, and standed at 4°C for 3h; Freeze-centrifuge at 4000rpm for 10 min, discarded the supernatant, added water to dissolve the protein precipitate, added Na_2_SO_4_ (the final mass fraction is 13%) to fully dissolve it, and then It was standed at 4°C for 3 h; Freeze-centrifuge at 4,000 rpm for 10min, discarded the supernatant, and add PBS for dissolution. Dialyzed the solution in a dialysis bag for 4-5 h and stored it at −20°C.

### 4. SDS-PAGE

A 10% SDS-PAGE separation gel was prepared according to the formula of the conventional process, followed by loading, electrophoresis, dyeing, decolorization, and gel imaging (Fig.3).

### 5. Western blot

SARS-CoV-2 S-RBD (provided by Sina Biological Company) was separated by polyacrylamide Amine gel (10% separation gel + 5% concentration gel) electrophoresis at 50 mA for 60 min. The protein on the gel was transferred to a NC membrane using the wet transfer method. The NC membrane was immersed in a 5% blocking solution, placed on a shaker, and blocked for 1.5 h at room temperature. After the membrane was washed three times for 5 min with TBST, SARS-CoV-2-IgY (dilution ratio: 1:10,000) was used as the primary antibody, and the primary antibody was incubated overnight using the inversion method. The membranes were then washed five times for 10 min each with TBST, immersed in blocking solution, and blocked for 30 min at room temperature. The secondary antibody was prepared by diluting HRP-labeled goat anti-chicken (provided by ZSGB-Bio) at a ratio of 1:10,000 and incubated for 1.5 h at room temperature, followed by five 5 min washing steps with TBST. The mixed immunoblotting chemiluminescence solution was dripped on the NC membrane, incubated at room temperature for 2-3 min, then placed in a cassette. The X-ray film was developed and scanned (Fig.4).

### 6. Cells and viruses

293T and Vero E6 cells were obtained from ATCC and maintained in DMEM (Gibco) supplemented with 10% fetal bovine serum. Prof. Ningshao Xia, Xiamen University, provided pseudotyped VSV-ΔG viruses expressing either a luciferase reporter or a mCherry reporter. The SARS-CoV-2 live virus (strain IVCAS 6.7512) was provided by the National Virus Resource, Wuhan Institute of Virology, Chinese Academy of Sciences.

The 293T, Vero E6 were obtained from ATCC and maintained in DMEM (Gibco) supplemented with 10% foetal bovine serum. The pseudotyped VSV-ΔG virusesexpressingluciferase reporter were provided by Prof. Ningshao Xia, Xiamen University. The SARS-CoV-2 live virus (strain IVCAS 6.7512) was provided by the National Virus Resource, Wuhan Institute of Virology, Chinese Academy of Sciences.

### 7. Pseudotype virus production

To produce pseudotyped VSV-ΔG-Luc bearing SARS-CoV-2 spike protein (pSARS-CoV-2), Vero E6 cells were seeded in 10 cm dish and transfected simultaneously with 15 μg SARS CoV-2-S-Δ18 plasmid by Lipofectamine 3000 (Thermo). Forty-eight hours posttransfection, 150µl pseudotyped VSV-ΔG bearing VSV-G protein were used to infect Vero E6 cells. Cell supernatants were collected after another 24 h clearing from cell debris by centrifugation at 3000rpm for 6 min, aliquoted and stored at −80°C.

### 8. SARS-CoV-2 entry assay based on pseudotyped virus

VeroE6 cells were seeded in 48-well plates and added 10ul volumes of pseudotyped VSV-ΔG-Luc bearing SARS-CoV-2 spike protein virus stocks with S-IgY or the control. At 24 h post-pseudotype-infection, the luciferase activities were measured with the Luciferase Assay System (Promega E4550).

### 9. qRT-PCR

According to the manufacturer’s instructions, 100 μL cell culture supernatant was harvested for viral RNA extraction using the MINIBEST Viral RNA/DNA Extraction Kit (Takara, Cat no. 9766). RNA was eluted in 30 μL RNase-free water. Reverse transcription was performed with a PrimeScript RT Reagent Kit with gDNA Eraser (Takara, Cat no. RR047A) and qRT-PCR was performed on a StepOnePlus Real-time PCR system (Applied Biosystems) with TB Green Premix Ex Taq II (Takara Cat no.RR820A). Briefly, 3 μLof total RNA was first digested with a gDNA eraser to remove contaminated DNA, and then the first-strand cDNA was synthesized in 20 μLreactions with 2 μL cDNA asthetemplate for quantitative PCR. The gene encoding the receptor-binding domain (RBD) was amplified by PCR from the cDNA template with primers: RBD-F: 5’-GCTCCATGGCCTAATATTACAAACTTGTGCC-3’; RBD-R: 5’-TGCTCTAGACTCAAGTGTCTGTGGATCAC-3’, cloned into pCMV-Flag vector (Invitrogen), and used as the plasmid standard after its identity was confirmed by sequencing. A standard curve was generated by determining copy numbers from serial dilutions (103–109 copies) of the plasmid. The primers used for quantitative PCR were RBD-qF1: 5’-CAATGGTTTAACAGGCACAGG-3’ and RBD-qR1: 5’-CTCAAGTGTCTGTGGATCACG-3’. PCR amplification was performed as follows: 95°C for 5 min followed by 40 cycles of 95°C for 15 s, 54 °C for 15 s, 72 °C for 30 s (Fig.5).

### 10. Immunofluorescence

To detect viral protein expression in Vero E6 cells, cells were fixed with 4% paraformaldehyde and permeabilized with 0.5% Triton X-100. Then the cells were blocked with 5% bovine serum albumin (BSA) at room temperature for 2 h. The cells were further incubated with the primary antibody (a monoclonal antibody against viral S protein) for 2 h, followed by incubation with the secondary antibody [Alexa 488-labeled goat anti-mouse(1:500; Abcam)]. The nuclei were stained with DAPI dye (Beyotime, China). The images were taken by fluorescence microscopy (Fig.6).

### 11. SARS-CoV-2 /VeroE6 cell experiment

25μL of SARS-CoV-2 (100PFU) was incubated with an equal volume of Pre-IgY, S-IgY and Remdesivir in different concentration gradients for 10 min, then added to 1×10^4^ Vero E6 cells, and cultured in an incubator at 37°C for 4 h, and then the cell culture solution was changed to complete medium. After 48 h, cells were collected and total RNA was extracted. The whole procedure was performed in a biosafety level (BSL-3) laboratory. After reverse transcription, the copy number of SARS-CoV-2 genome in each sample was detected by qRT-PCR(Fig.7).

## Author Contributions

Jingchen Wei and Yunfei Lu contributed equally to this work.

## Acknowledgements

This work was supported by the National Natural Science Foundation of China (No. 81560574). The work was also funded by Guangxi Liver Injury and Repair Molecular Medicine Collaborative Innovation Center, Guilin Medical University and The Guangxi Bagui Scholars Program to Songqing He.

We thank LetPub (www.letpub.com) for its linguistic assistance and scientific consultation during the preparation of this manuscript.

## Statistical analysis

Statistical analysis was performed using GraphPad Prism 6 software. Data are expressed as mean ± SD. T-test was used for two-group comparisons. The ^*^*P*< 0.05, ^**^*P*< 0.01, ^***^*P*< 0.001 were considered significant. Unless otherwise noted, error bars are indicated as mean values with standard deviation of at least three experiments.

